# Recurrent genetic abnormalities in human pluripotent stem cells: definition and routine detection in culture supernatant by targeted droplet digital PCR

**DOI:** 10.1101/658484

**Authors:** Said Assou, Nicolas Girault, Mathilde Plinet, Julien Bouckenheimer, Caroline Sansac, Marion Combe, Joffrey Mianné, Chloé Bourguignon, Mathieu Fieldes, Engi Ahmed, Thérèse Commes, Anthony Boureux, Jean-Marc Lemaître, John De Vos

## Abstract

Genomic integrity of human pluripotent stem cells (hPSCs) is essential for research and clinical applications. However, genetic abnormalities can accumulate during hPSC generation and routine culture and following gene editing. Their occurrence should be routinely monitored, but the current assays to assess hPSC genomic integrity are not fully suitable for suchregular screening. To address this issue, we first carried out a large meta-analysis of all hPSC genetic abnormalities reported in more than 100 publications and identified 738 recurrent genetic abnormalities (i.e., overlapping abnormalities found in at least five distinct scientific publications). We then developed a test based on the droplet digital PCR technology that can potentially detect 94.3% of these hPSC recurrent genetic abnormalities in DNA extracted from culture supernatant samples. This test can be used to routinely screen genomic integrity in hPSCs.

## Introduction

Human pluripotent stem cells (hPSCs) are stem cells that endlessly self-renew *in vitro* and that can differentiate into all adult cell types. Therefore, they are a potentially infinite and physiologically relevant cell material for research (*in vitro* modeling of human development and diseases) and regenerative medicine/cell therapies. These cells are isolated from discarded human embryos (i.e., human embryonic stem cells; hESCs), or obtained from differentiated cells by cell reprogramming (i.e., human induced pluripotent stem cells; hiPSCs). It is crucial that hPSC genome remains the faithful genetic copy of the cells from which they were derived. However, genetic abnormalities (e.g., karyotype abnormalities) can arise in hPSCs, for instance during cell reprogramming, cell culture, or genome editing (Assou et al., 2018). Many of these genetic abnormalities are often recurrent. For instance, gains of chromosome 12 (most frequently 12p), 17 (particularly 17q), 20 or X have been often detected using standard cytogenetic procedures (G-banding) (Lefort et al., 2009). Sub-chromosomal abnormalities, such as 20q11.21 amplification, also can be recurrent. The biological significance of such recurrent abnormalities is still discussed, but they might result in a strong selective growth advantage for cultured cells, as already demonstrated for the 20q11.21 amplification (Zhang et al., 2019). Therefore, it is crucial to carefully catalogue all genetic alterations found in hPSCs and identify the recurrent ones. To this aim, we carried out a meta-analysis of published genetic abnormalities found in hPSCs. We could give a precise definition of recurrent genetic abnormality and then listed all of them in a large dataset. As these recurrent genetic abnormalities are found in specific genomic regions, we developed a focused droplet digital PCR (ddPCR) approach that allowsscreening more than 90% of these recurrent abnormalities in DNA isolated from cell culture supernatant. This method greatly simplifies and therefore encouragesthe regular and systematic hPSCs screening.

## Results

### Meta-analysis of hPSC genetic abnormalities and identification of a recurrencepattern

To catalogue all genetic abnormalities previously detected in hPSCs using various techniques (karyotyping, fluorescence in situ hybridization, comparative genomic hybridization, microarray-based comparative genomic hybridization, and next generationsequencing, NGS), we selected primary research articlesthat reported genetic abnormalities in hESCs and hiPSCs, and extracted the DNA abnormalitygenomic coordinates as well as the experimental data to characterize these abnormalities. We collected data on 942 cell samples and on 415 750 variants and abnormalities from 104 different studies published between 2004 and 2016 (Figure 1A-B, and Table S1). The dataset included the major publications on genetic abnormalities in hPSCs during culture and also articles that identified one or several abnormal clones in new hPSC lines. A first global analysisof all listed mutations allowed identifying genome locations where these abnormalities were more frequently localized: trisomy 12 and 12p amplification, 20q11.21 amplification, trisomy 17 and 17q amplification, chromosome 1 amplification, and trisomy X (female cell lines) (Figure S1A). Abnormalities that accumulated at a specific genome location (i.e., recurrent abnormalities) weremostly aneuploidy or copy number variations (CNV), in agreement with previous reports. No abnormality smaller than 10 base pairs (bp) displayed a recurrent profile in this large dataset (Figure S1B). We also reported 93 translocations and 20 inversions, involving all chromosomes with the exception of chromosome 12, mainly of chromosomes 1 and 17, but without a clear recurrent pattern (Figure 1C). As described previously (Bai et al., 2015), genetic abnormalities were more frequently reported when hPSCs were cultured using enzymatic passaging compared with mechanical passaging (Figure 1D-E). However, the frequency of abnormalities associated with enzymatic passaging decreased after 15 passages.

In summary, this large meta-analysis of genetic abnormalities in hPSCs confirms the recurrence of large CNVs and chromosomal abnormalities, and provides a large dataset of recurrent abnormalities.

### Definition and analysis of recurrent genetic abnormalities

As no quantitative definition of a recurrent hPSC genetic abnormality exists in the literature, we wanted to establish a clear threshold for such events. A recurrent genetic abnormality was therefore defined as a non-polymorphic variant that overlaps with abnormalities found in other hPSC lines. The recurrence pattern most likely reflects a common functional cause that occurs in different laboratories and in different cell lines (Assou et al., 2018). We hypothesized that the abnormalities with the strongest functional impact on hPSC growth would be those that are (1) common to different hPSC lines and (2) found in different culture conditions. We estimated that a genetic abnormality met these two criteria if all/part of the altered sequence overlapped with that of other genetic abnormalities that weredescribed in at least four other distinct scientific publications (thus, at least five articles in total) (Figure 2A).To identify recurrent genetic abnormalities based on thesecriteria, we analyzed all variants>10bp that were not polymorphisms (n = 8284). We found that recurrent abnormalities are only CNVs (including chromosomal gain or losses) (Figure 2B). By plotting the genomic coordinates of these 738 recurrent abnormalities, we found that they were mainly localized in known hotspots, such as chromosome 1, 12, 17q, 20q11.21 or X (Figure 2C). Conversely, there were no recurrent abnormalities in chromosome 2, 4, 10 or 21.

**Figure 1.**
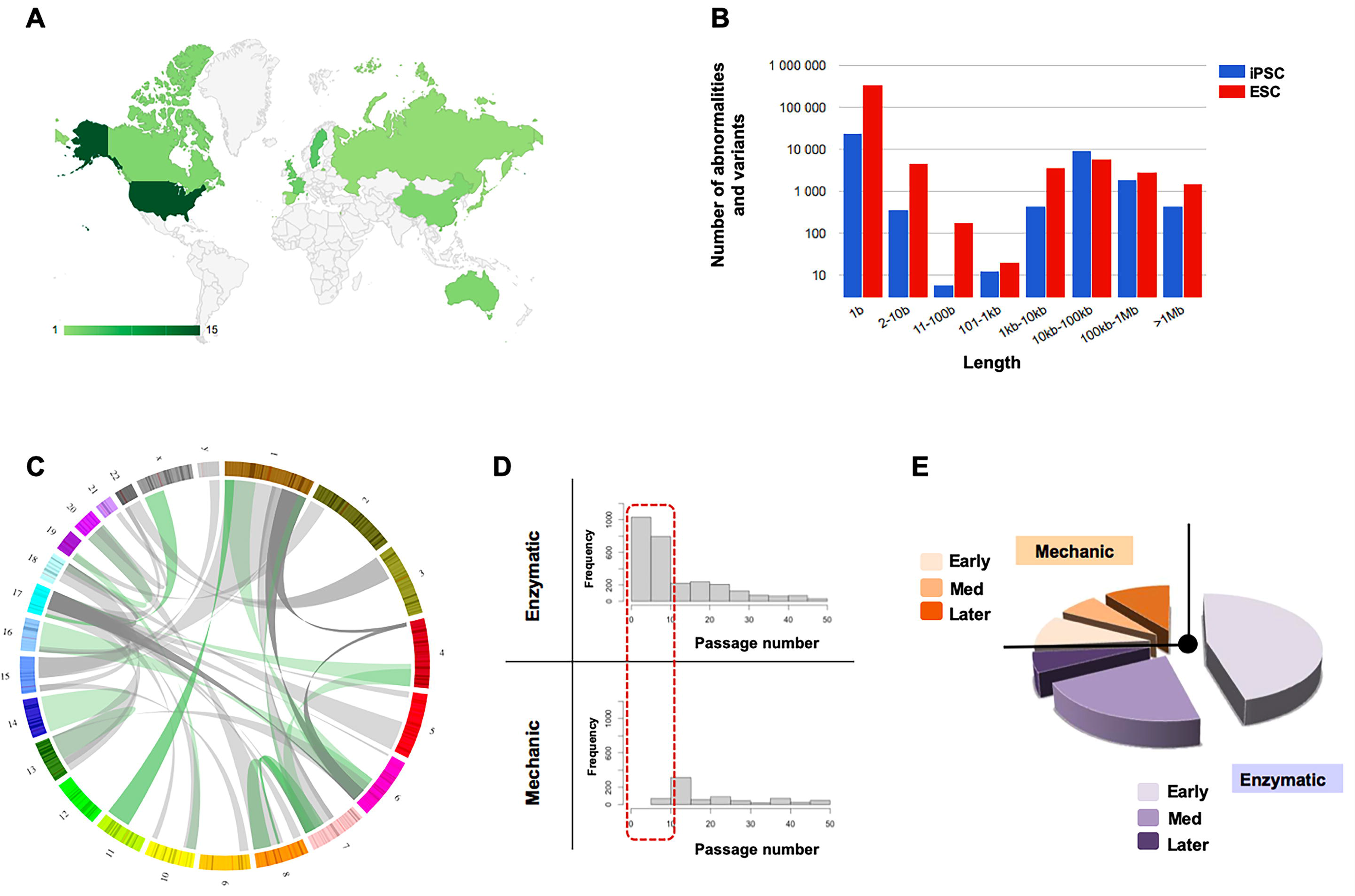
**A.** Countries contributing to the articles included in the analysis. **B.** Number of genetic abnormalities and variations collected in this study, according to their length. **C.** Circos plot representing all translocations reported in the articles analyzed in this study. Numbers indicate chromosome number. Balanced translocations are in green, unbalanced translocations in grey. **D.** Frequency of genetic abnormalities and variantsfor which information on previous passage technique was available, according to the passage number and passaging technique (mechanic vs enzymatic). **E.** The genetic abnormalities were grouped according to the hPSC passage number into: early (1 – 15 passages; Early), intermediate (15 – 50 passages; Med) and later (> 50 passages) passage, and according to the passaging technique (mechanic vs enzymatic).

We then investigated the nature of these hotspot regions (Figure 2A) (Table 1, lists the ten more frequent common regions). Specifically, four regions included more than 50% of all reported abnormal genetic abnormalities. Moreover, a limited set of common abnormal regions comprisedthe most recurrent genetic abnormalities. Indeed, more than 95% of recurrent abnormalities were restricted to 24 common regions. A set of probes designed to cover these regions would detect all genomic abnormalities of these regions, except balanced translocation, i.e. 94.3% of all recurrent genetic abnormalities (Figure 2D).

**Figure 2.**
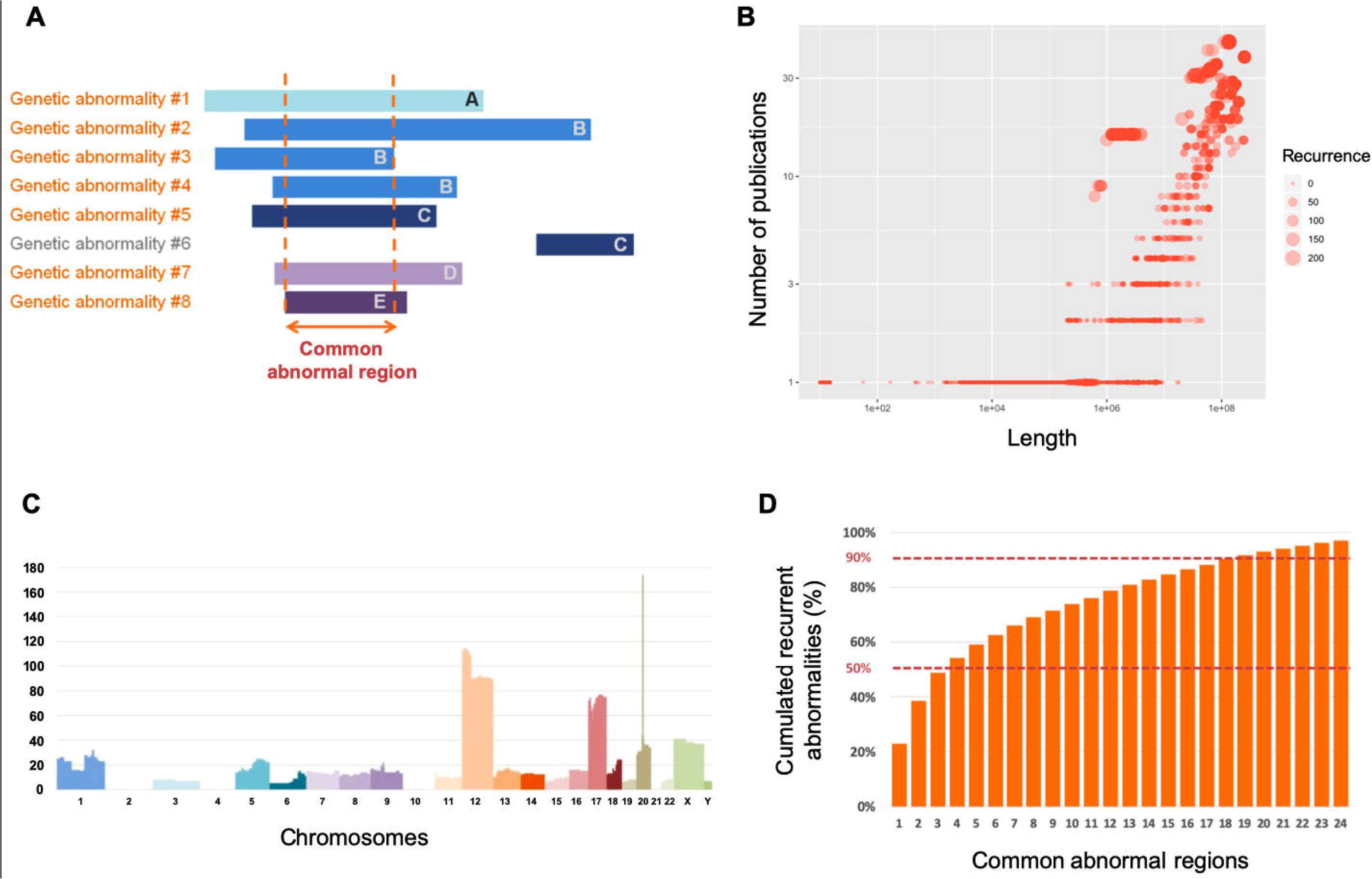
**A.** Graphic representation of eight genetic abnormalities (or variants that are not polymorphic) (#1 to #8), from five different articles (A to E, one color for each article). Part or the entire sequence of seven abnormalities (in orange) overlapped and were classified as recurrent genetic abnormalities because they were from five different articles. Conversely, abnormality #6 is not recurrent. The region shared by the seven recurrent genomic abnormalities is a common abnormal region. **B.** Dot plot showing all genetic abnormalities larger than 10bp and that were not polymorphisms (n=8 284). Abnormalities are plotted according to their length (xaxis), the number of other abnormalities with which they overlapped (recurrence: diameter of the dots) and the number of different articles that described these overlapping abnormalities (y axis). Recurrent genomic abnormalities (≥5 different articles) are above the dashed violet line. **C.** Bar plot showing the 738 recurrent genetic abnormalities according to their genomic coordinates (one color per chromosome). Y axis: number of recurrent abnormalities. **D.** Percentage of cumulated recurrent abnormalities found in 24 common abnormal regions comprising the most recurrent genetic abnormalities.

**Table 1:** Common abnormal regions. The 10 most common abnormal regions ranked according to the number (n) of abnormalities that include that region. Percentage: number of recurrent abnormalities that share a common abnormal region relative to all recurrent abnormalities (Reference genome: GRCh37/hg19).

### Cell culture supernatant as a DNA source to evaluate hPSC genomic integrity by ddPCR

We then decided to take advantage of this highly biased recurrence profile of hPSC genetic abnormalities to develop a rapid PCR-based approach to detect the most common recurrent abnormalities, including those that cannot be detected by karyotyping due to its resolution limits. We analyzed DNA extracted from different hPSC lines (cell-DNA) without (HY03, UHOMi001-A, iCOPD2 and iCOPD9) and with genetic abnormalities (RSP4: chromosome 20 triploidy), using ddPCR and primer pairs that target chromosome X or chromosome 20. We could observe two chromosome 20 copies in DNA samples from normal hPSC lines, and three copies in the RSP4 cell-DNA sample (Figure 3A and S2). Second, to test the sensitivity of our approach, we prepared cell-DNA from UHOMi001-A cells (diploid line) mixed with increasing percentages (0% to 100%) of HD291 cells (chromosome 12q trisomy). Our ddPCR approach could detect the presence of the chromosome abnormality, starting from the sample containing 10% of HD291 cells (Figure 3B).

**Figure 3.**
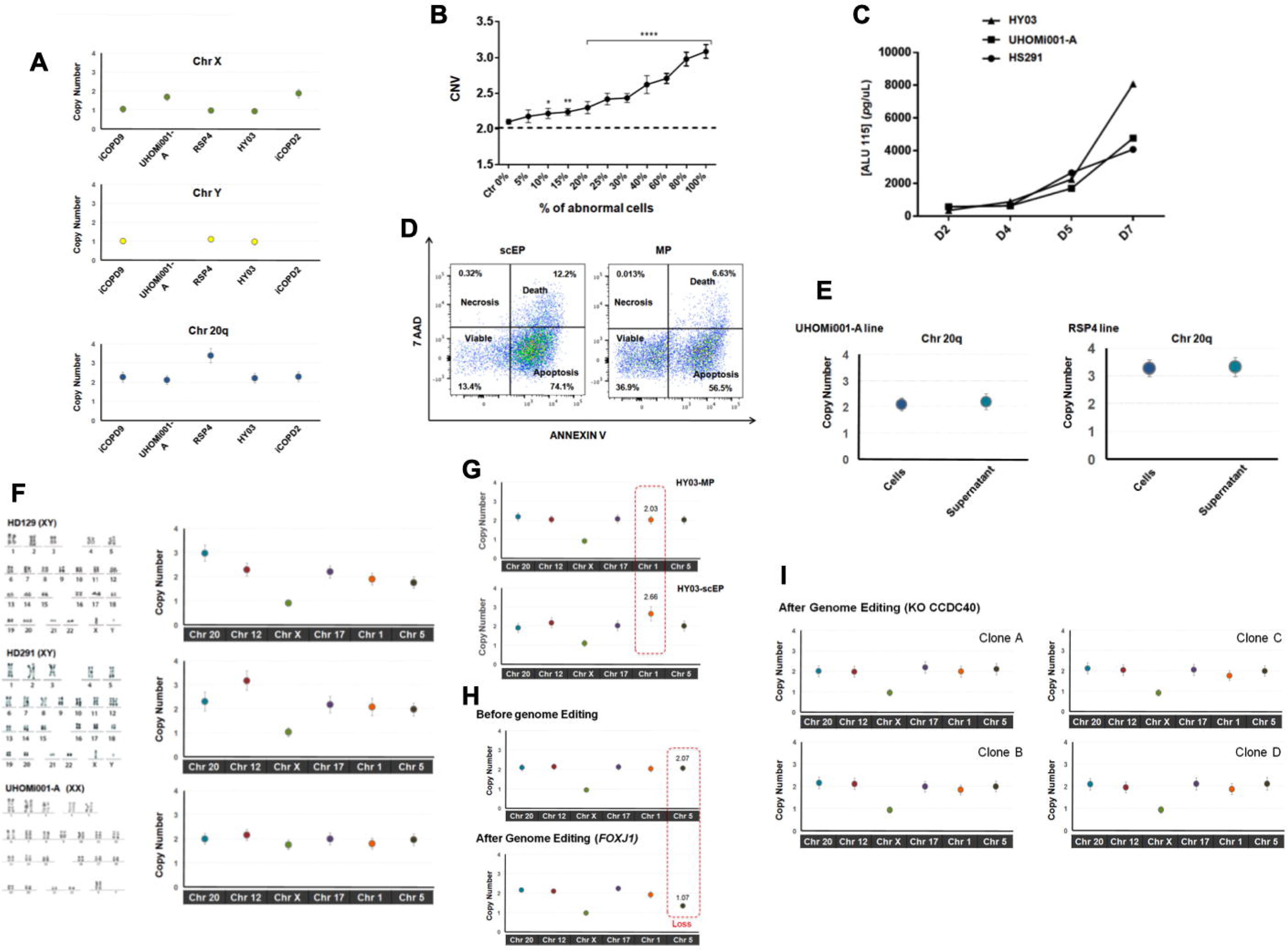
**A.** Copy number variation analysis using droplet digital PCR and DNA extracted from different hPSC lines in culture. Error bars indicate the Poisson distribution at 95% confidence intervals. **B.** Sensitivity of the droplet digital PCR method for detecting increasing percentages (from 0 to 100%) of hPSCs harboring a trisomy 12 within a sample of euploid hPSCs. A significant difference in the CNV compared with control (0%) was observed in samples with at least 10% of abnormal cells (p <0.001). The panels represent three biological replicates. All values of copy number were corrected by adding −0.083 as more than 50 experiments on samples with a normal count of chromosome 12 have shown a bias with this probe with a median value of 2.083. **C.** Quantification of cell-free DNA in supernatant samples from one hESC line (HS291) and two hiPS lines (names HY03; UHOMi001 cells) cultured in E8 medium on Geltrex matrix. Supernatant was collected at the indicated time points after seeding (75,000 cells/well of a 35mm plate). DNA was extracted from 300 μL of supernatant and quantified by ALU-qPCR with ALU115 primers. **D.** The percentage of apoptotic, necrotic and viable cells in supernatant samples collected at day 5 was evaluated by flow cytometric analysis after staining with Annexin-V and 7-Amino-Actinomycin (7-AAD). Every dot corresponds to a single cell. scEP: single-cell Enzymatic passaging; MP: Mechanical passaging. **E.** Copy number of chromosome 20q measured by ddPCR using genomic DNA from cells and supernatant as template. The error bar varies in function of the DNA source (cells or supernatant). **F.** The iCS-digital test using six probes corresponding to the abnormal genomic regions with the highest overlapping recurrent genomic abnormalities can identify aneuploidy. The hPSC lines HD129 (chromosome 20 triploidy), HD291 (chromosome 12 triploidy) and UHOMi001-A (euploid) were analyzed by karyotyping (classical G-banding) and with the iCS-digital test using six specific probes targeting the common abnormal regions onchromosomes 20q, 12, X,17, 1 and 5. Karyotype image for HD129 and HD291 hESCs lines are from (Bai et al., 2015) with permission from the publisher (NB: this permission should be requested to Stem Cell Dev). **G.** Identification of genome modifications associated with single-cell enzymatic passaging using the iCS-digital test. The HY03 hiPS line remained euploid using mechanical passaging (MP), whereas after 15 passages using single-cell enzymatic passaging (SC), it displayed a copy number gain on chromosome 1. **H.** Analysis of genome stability using the iCS-digital test in hiPSC line HY03 (male cells) before and after genome editing revealeda 5q loss in FOXJ1 mCherry edited cells. CCDC40 knock-out (KO) clones obtained after CRISPR/Cas9 editing remained euploid.

Another major constraint to hPSC genome integrity analysis is the need to dedicate a significant part of the cell culturefor this purpose. Therefore, we investigated whether genome integrity could be assessed using DNA extracted from the hPSCculture supernatant. We found that when using supernatant-derived DNA (supernatant-DNA), the concentration of DNA fragments, estimated by quantitative PCR (qPCR), increased progressively during hPSCs culture and was at a highest after 7 days in culture, when cell confluence was higher than 70% (Figure 3C). To helpelucidating the origin of the supernatant-DNA, we analyzed floating cells by staining with Annexin V and 7ADD followed by flow cytometry (Figure 3D). This analysis showed that 74.5% and 56.5% of floating cells were apoptotic, 12.2% and 6.6% were dead, and 13.4% and 36.9% were viable after single-cell and mechanical passaging respectively. We also measured the DNA integrity index by qPCR, using the ALU115 primers that amplify both short (apoptotic) and long (non-apoptotic) DNA fragments, and the ALU247 primers that amplify only long non-apoptotic DNA fragments (Umetani et al., 2006). The ALU115/ALU247 ratio was higher than two, showing that supernatant contained a majority of apoptotic cells (Figure S3). We also investigated the influence oftime delay before DNA extraction and freeze-thaw cycles number on the quantity of DNA extracted from cell culture supernatant samples (Figure S3). These results demonstrated that stable and significant quantities of DNA can be extracted from cell culture supernatant samples.

Then, we observed an excellent agreement between the results obtained using cell-DNA and the corresponding supernatant-DNA (supernatant collected at day 5-7 of culture) (Figure 3E). This shows that ddPCR offers the sensitivity required for evaluating genomic integrity insupernatant-DNA, and that supernatant-DNA could be used instead of cell-DNA.

We then evaluated whether supernatant-DNA could be used to perform focused ddPCR on culture supernatant (in culture supernatant digital PCR test, iCS-digital test) using a panel of six commercial pre-designed probes that correspond to or are close to the sixmost common abnormal regions (chromosomes20q, 12, 17, X, 1 and 5). These six probes target genome regions that comprise more than 50% of all recurrent genetic abnormalities found in hPSCs (61% cumulated coverage of recurrent abnormalities).We analyzed supernatant-DNA from two hPSC lines with abnormal karyotype (HD129 and HD291: chromosome 20 and 12 triploidy, respectively) (Bai et al., 2015) and one diploid line (UHOMi001-A). The iCS-digital test results overlapped with those obtained by karyotyping (Figure 3F). In conclusion, targeted ddPCR can efficiently detect CNVs and can be carried out using supernatant-DNA.

### Routine screening of hPSClines during cell culture and after CRISPR gene editing using the iCS-digital test

The simplicity of the iCS-digital test could allow the routine screening of the most recurrent genetic abnormalities in hPSC lines, particularly when using single-cell or small-clump passaging, a major cause of genomic alterations (Bai et al., 2015) (Figure 1F-G). The iCS-digital test revealed that theHY03hiPSC line, which was euploid at passage 5, harbored a chromosome 1 gain at passage 15after single-cell passaging, but not after mechanical passaging (Figure 3G).

Cell reprogramming and genome editing using CRISPR/Cas9 require clonal expansion/selection that favors the emergence of abnormal clones. The iCS-digital test indicated no alteration at chromosomes 20q, 12, 17, X, 1 and 5 in the hiPSC line HY03 (passage M53Cl2SC11) before genome editing (Figure 3H). Conversely, after introduction of them Cherry cassette at the 3’ of the *FOXJ1* gene, or knock-out (KO) of the CCDC40gene using the CRISPR/Cas9 methodology, the iCS-digital test showed that the single clone identified to be correctly edited (passage 7) harbored a CNV of the long arm of chromosome 5 (copy number=1.3), while the four CCDC40_KO clones analyzed appeared euploid for the six regions checked (Figure 3I). Taken together, our results show that focused ddPCR can be used to rapidly screen iPSCs after derivation, during cell culture or amplification, and after cell cloning in settings such as gene editing.

## Discussion

The present study is, to our knowledge, the largest meta-analysis of hPSC genetic abnormalities in more than 100 different research articles from many different laboratories and cell lines. This allowed us to propose a quantitative threshold to define recurrent genetic abnormalities in hPSCs and to test this threshold. Hence, a recurrent genetic abnormality is an abnormality that shares part of its abnormal sequence with other abnormalities that have been reported in at least five different publications. Our threshold should favor the detection of abnormalities that are common to different hPSC lines and different laboratories, as opposed to abnormalities that are specific to a unique cell line, a hiPSC donor cell or to the culture conditions used in one laboratory. Our analysis confirmed that CNVs are the main recurrent genetic abnormality in hPSC lines. In agreement, a recent extensive survey of NGS data to identify recurrent small mutations reported that only TP53 was prone to recurrent mutations in hPSCs (14 mutated samples among the 257 independent hPSC lines studied, 5.45%) (Merkle et al., 2017). It has been proposed that the high hPSC susceptibility to mitotic division errors and to the upregulation of anti-apoptotic proteins contribute to the high frequency of aneuploidy and CNVs at specific genomic locations (Zhang et al., 2019). Indeed, we and othersidentified a limited list of genome regions that group overlapping recurrent genomic hPSC abnormalities, possibly because these changes provide a growth advantage to cultured cells. For instance, chromosome 20q11.21 gain leads to upregulation of the anti-apoptotic protein BCL-XL (Zhang et al., 2019). Importantly, this biased distribution makes possible to study common abnormal regions by targeted PCR. For instance, by targeting only the four most common abnormal regions, more than 50% of all recurrent genetic abnormalities are covered, and more than 90% by targeting 24 regions. Screening of recurrent genetic abnormalities is paramount to claim that hPSCs are normal (Bai et al., 2013; International Stem Cell Initiative et al., 2011). Although a targeted approach is by definition not exhaustive, it is an effective strategy to rule out the most frequent and functionally damaging abnormalities found in hPSC lines.

We demonstrated that this strategy can be carried out simply by usingDNA extracted from culture supernatant. This DNA reliably reflected the DNA profile of adherent cells. Moreover, ddPCR is a robust and sensitive technology to assess CNV on targeted abnormal genomic regions to screen the most common recurrent abnormalities.

Genomic integrity of hPSCs in culture should be frequently assessed. We recently noted in a series of 25 consecutive studies on hiPSCs that the current genomic screening practices were unsatisfactory because no genomic integrity follow-up was carried out for any of the hiPSC lines (Assou et al., 2018). This could be explained by the labor and costsinvolved in the implementation of classical screening techniques, such as karyotyping. Therefore, a simple test that canrapidly rule out the most frequent recurrent genomic abnormalities might promoteadhesion to good practices for hPSC genomic integrity screening. Moreover, karyotyping can miss abnormalities that are smaller than 5 – 10 Mb. For instance, we found that among all 170 recurrent genomic abnormalities on chromosome 20, 168 overlapped with 20q11.21 and among them 135 were smaller than 5 Mb. By contrast, the probes of the iCS-digital test can detect all 168 20q11.21 abnormalities.

In conclusion, we used a large dataset of hPSC genomic abnormalities based on more than 100 publications to strictly define recurrent genetic abnormalities. Our exhaustive database of such genomic defects allowed identifying a set of common abnormal genomic regions that involve more than 90% of all recurrent abnormalities. This offered the opportunity to develop and evaluate the efficacy of a targeted ddPCR approach. Moreover, we show that culture supernatant contains enough DNA to perform ddPCR. Therefore, we propose a simple test based on supernatant DNA (iCS-digital test) that could be used to routinely screencultured hPSC lines.

## Experimental Procedures

### Analyzing recurrence

All genetic abnormalities were converted using their GRCh37A genome coordinates. A first analysis was carried out using a recurrence score (RS) for data split in two datasets (>10 bp and ≤10 bp) from which polymorphic data (sequences present in dbSNP or DGV) were removed. The RS was computed by comparing each abnormality to all the others and by identifying abnormalities with a reciprocal overlap of at least 0.2. Regions with a reciprocal overlap higher than 80% were merged. For each overlap, RS was computed as follows: RS = a * s, where (a) is thenumber of abnormalitiesthat contributed to define this overlap (identicalabnormalities from the same cell line in the same study were counted only once) and (s) the number of different studies from which these overlapping abnormalities came from.

Then, to define recurrent genetic abnormalities, each abnormality >10 bp and that was not a polymorphism (n=8284 abnormalities in total) was compared with all the others, and abnormalities with a reciprocal overlap >0.33 and larger than 0,2kb, and the number of publications from which these abnormalities originated were counted. Abnormalities that overlapped with other abnormalities that came from at least four other publications (number of total publications ≥5) were defined as recurrent and kept, and this process was carried out iteratively and rapidly converged on a stable list of 738 recurrent abnormalities. Loss and gain were considered indiscriminately because the aim was to identify common abnormal regions. Bedtools was used to identify common abnormal regions.

### Cell culture

The human hESC lines HD129 and HD291 were derived in our laboratory (Bai et al., 2015; Ramirez et al., 2011). The hiPSC lines UHOMi001-A (Ahmed et al., 2018), RSP4, iCOPD2A1, iCOPD9A2, and HY03 were reprogrammed using the Sendai virus and the CytoTune^®^-iPS 2.0 Sendai Reprogramming Kit (Thermo Fisher Scientific), and display all the PSC features: grow as typical PSCs, are positive for pluripotency markers (*OCT4, NANOG, SOX2, TRA1-60, TRA1-81, SSEA3, SSEA4*) and for phosphatase alkaline activity, and can differentiate into cells of all three germ layers. PSC lines were maintained on Geltrex matrix (Thermo Fisher Scientific) in Essential 8 (E8) Medium (Thermo Fisher Scientific) or on Matrigel (BD Biosciences) in mTeSR1™ medium (Stem Cell Technologies), according to the manufacturer’s instruction. Cells were grown in 35-mm dishes at 37□°C and were passaged either mechanically or dissociated into single cells or small clumps every week. Mechanical passaging (MP) was carried out under an inverted microscope in a hood using scalpels. For single-cell enzymatic passaging (scEP), clump passaging, colonies were pre-incubated with the Rho-associated protein kinase (ROCK) inhibitor Y-27632 for 1h, and then dissociated with TrypLE™ Select (Invitrogen) or EDTA (Versene Solution, Thermo Fisher Scientific) at 37°C for 10 min.

### Culture medium collection and nucleic acid extraction

Supernatant (1.5 mL) was collected into a safe-lock tube (DNase-free) before passaging from cell cultures that were at least 70% confluent. DNA was extracted from 200 μL of supernatant using the QIAmp DNA Mini Blood Kit (Qiagen, Hilden, Germany) according to the manufacturer’s protocol. Briefly, 20 μL proteinase K and 200 μL Buffer AL were added to each supernatant sample. After pulse vortexing for 15s, the lysis mixture was incubated in an Eppendorf tube (1.5 mL) at 56°C for 10min. The highly denaturing conditions and elevated temperatures favored the complete release of DNA from any bound proteins. After adding 200 μL cold ethanol (100%) to the lysates, samples were transferred in QIAamp Mini columns and centrifuged at 6000g for 1min followed by two wash steps (in Buffer AW1 and Buffer AW2) to eliminate contaminants. Then, supernatant-DNA was eluted in 60 μL Buffer AE and stored at −20°C.

### Quantification of supernatant-DNA by ALU-qPCR and QuBit

Supernatant-DNA was analyzed by qPCR (LC480, Roche) using the ALU 115 and ALU 247 primers, as previously described in (Umetani et al., 2006). One μl of each eluted supernatant-DNA sample was added to a reaction mixture containing 2X Light Cycler480 SYBR Green I master mix (Roche Applied Science, Germany) and 0.25 μM of forward and reverse primers (ALU-115 and ALU-247) as described in (Umetani et al., 2006) in a total volume of 10□ μL. Reactions were carried out in 96-well white plates using an EpMotion 5070 Liquid Handling Workstation (Eppendorf). All reactions were performed in triplicate. A negative control (RNAse/DNAse-free water) was included in each run. The supernatant-DNA concentration was determined using a standard curve obtained by successive dilutions of a commercial human genomic DNA sample. DNA integrity was calculated as the ratio of the qPCR results with the two primer sets (ALU115 and ALU247). The ALU115 set amplifies smaller fragments that result from apoptosis and the ALU247 set amplifies only larger fragments that result from necrosis. Supernatant-DNA concentration was quantified using the QuBit dsDNA HS Assay Kit and aQuBit 2.0 fluorometer following the manufacturer’s instructions (Life Technologies).

### Flow cytometric detection of apoptosis and necrosis using the Annexin V and 7ADD assay

The PE Annexin V Apoptosis Detection Kit I (BD phramingen, Ref: 559763) was used to quantify the percentage of apoptotic and necrotic cells in supernatant samples. Briefly, samples were incubated with PE Annexin V in a buffer containing 7-Amino-Actinomycin (7-AAD) according to the kit protocol (http://www.bdbiosciences.com/ds/pm/tds/559763.pdf), and analyzed by flow cytometryat the Montpellier Resources Imaging (MRI) facility (https://www.mri.cnrs.fr/en).

### Digital droplet PCR (ddPCR)

The ddPCR workflow was performed according the Bio-Rad instructions (Bio-Rad QX200 system). Briefly, reactions were set up using one primer pair that targets the region of interest (for instance: CNV-chr20) and a second primer pair that targets the reference gene (*RPP30*). The two primer sets are labeled with different fluorophores (FAM and HEX). DNA (amount) from each samples was added to the TaqMan PCR reaction mixture (final volume of 20 μL) that included 2X Supermix No dUTP (Bio-Rad, Ref: 1863023) and the primer sets. Each reaction mixture was loaded in a disposable plastic cartridge (Bio-Rad) with 70 μL of droplet generation oil (Bio-Rad) and placed in the droplet generator (Bio-Rad). The cartridge was removed from the droplet generator, and the droplets collected in the droplet well were then manually transferred with a multichannel pipette to a 96-well PCR plate. The PCR amplification conditions were: 94°C for 10min, 40 cycles of 94°C for 30s, and 60°C for 1min, followed by 98°C for 10min and ending at 4°C. After amplification, the plate was loaded into the QX200 Droplet Reader (Bio-Rad). Copy number was assessed using the Quantasoft software. For testing the ddPCR sensitivity in detecting a gain of CNV-12q, the UHOMi001-A diploid and the HD291 aneuploid line were used. Cells were grown on Geltrex matrix in E8 Medium prior to the experiment and then dissociated using trypsin and counted. After mixing the two cell lines to obtain increasing concentrations (from 0% to 100%) of abnormal cells within the diploid population, each mixed sample was processed for genomic DNA extraction using the QIAmp DNA Mini Blood Kit (Qiagen, Hilden, Germany) and for ddPCR analysis. Reference for the designed BioRad ddPCR probes are the following: chromosome 20 probe #dHsaCP2506319; chromosome 12 probe #dHsaCP1000374; chromosome X probe #dHsaCP2506654; chromosome 17 probe # dHsaCP1000054; chromosome 1 probe #dHsaCP1000482; chromosome 5 probe # dHsaCNS501868922 and RPP30 probe # dHsaCP2500350.

### Generation of the FOXJ1_mCherry and CCDC40_KO iPSC lines using CRISPR/Cas9

A stock of HY03 (75k) M53Cl2SC6 cells was made and their euploidy after thawing was confirmed using the iCS-digital test for detection of CNV anomalies at M53Cl2SC11 or M53Cl2SC17 for FOXJ1_mCherry tagging and CCDC40_KO projects respectively. The day before transfection, 25000 HY03 (75k) M53Cl2SC7 cells per cm^2^ were plated in a 6-well plate coated with Geltrex matrix and with E8 Medium supplemented with Y-27632 (10μM). The day of transfection, medium was refreshed at least 2 hours before transfection using the Lipofectamine Stem transfection reagent (Invitrogen), following the manufacturer’s instructions. Briefly, 2 μg of pSpCas9(BB)-2A-GFP (PX458) (gift from Feng Zhang, Addgene plasmid #48138) containing the FOXJ1 targeting sgRNA sequence 5’-GGGCCTTCTTGTAAGAGGCC-3’ or the CCDC40 targeting sgRNA sequence 5’-CTCCTCGTTGGCGGCTGCGCAGG-3’ with 1 μg of MBX plasmid (gift from Linzhao Cheng, Addgene plasmid #64122) and 1 μg of homemade donor plasmid pUC19_FOXJ1_mCherry_cNEO for the FOXJ1_mCherry tagging project were mixed with 4 μL of Lipofectamine and left at room temperature for 10 min to form complexes. The Lipofectamine-DNA complexes were added on top of the cells, distributed by gently swirling the plate, and incubated at 37°C, 5% CO_2_. The following day, the medium was changed with fresh E8 medium supplemented with G418 (200 μg/ml) for the FOXJ1_mCherry project), media was changed daily for 6 days and colonies were manually picked and transferred into a well of a 96 well plate for amplification. At confluence, clones were passaged to a 24-well plate, and then to a 6-well plate. DNA was collected to screen clones by bridge-PCR, transgene copy counting and Sanger sequencing for FOXJ1_mCherry tagging or by high resolution melt analysis (HRMA) followed by Sanger sequencing for CCDC40_KO. Finally, the presence of genomic abnormalities was checked using the iCS-digital test.

### Statistical Analysis

For ddPCR, absolute quantification was based on the number of positive droplets and Poisson sampling statistics, as follows: λ = −ln(1-k/n) where k is the number of positive droplets, n the total number of droplets and λ the mean copies per droplet. To test the statistical difference of the ddPCRresults between samples with different percentages of trisomic cells, first values were normalized to those of the 0% sample within each biological replicate. Data were analyzed using Excel. p <0.05 defined statistical significance.

## Supporting information

Bubble plot showing recurrent DNA abnormalities

FACS-like plot illustrating ddPCR results

Effects of various preanalytical conditions on supernatant DNA

Articles analyzed to identify genetic abnormalities in hPSC lines

## Author Contributions

S.A., N.G., and J.D.V designed the study and analyzed data; S.A., M.C., J.M., E.A., performed the experiments; M.P., J.B., C.S., C.B., M.F., E.A., collected and analyzed data; T.C., A.B. analyzed and interpreted the data; S.A., and J.D.V wrote the paper. J.M.L revised the manuscript. All authors approved the final version prior to submission.

## Acknowledgements

We thank Camille Novo for contributing to the collection of PSC genomic abnormalities, Elena Hauser for technical help and Guilhem Requirand for his expertise in flow cytometry. This study was funded by ÌNSERM, ANR (INGESTEM) and the SATT AxLR.

## Supplemental Information

### Supplemental figure legends

**Figure S1**

**A.** Bubble plot showing recurrent DNA abnormalities >10bp in hPSClines. Each bubble represents a region where several DNA abnormalities from several publications are found. The bubble size is proportional to the region length and the color indicates the number of publications that reported abnormalities in that region. The horizontal axis corresponds to the genome position and the vertical axis corresponds to the recurrence score (see Material and Methods).

**B.** Same as in C, but for abnormalities and variants ≤10bp.

**Figure S2**

**A.** FACS-like plot showing a typical result obtained from quantifying chromosome 20q CNVin a euploid hPSCline (UHOMi001-A) using DNA from cells and supernatant (Sup.), as indicated. Blue: droplets positive for the target CNV (Chr20q); green: droplets positive for the reference gene (*RPP30*); grey: negative droplets (containing no target or reference genes); orange: droplets positive for both the reference and target genes.

**B.** FACS-like plot showing a typical result obtained from quantifying chromosome 20q CNVin an aneuploid hPSCline (RSP4) using DNA from cells and supernatant (Sup.), as indicated. Same color code as in A.

**Figure S3**

**A. Concentration of DNA in supernatant samples and effects of various preanalytical conditions.** Quantification of supernatant-DNA by quantitative PCRwith two sets of ALU primers (115 and 247 bp) that amplify DNA fragments of different length in supernatant samples collected from three hPSC lines at day 5 and day 7. The ALU 115 and ALU 247 values are significantly different (p<0.05). The Q247/Q115 ratio indicates the DNA integrity value (Q115 corresponds to the DNA concentration obtained using the ALU 115 primers and Q247 to the concentration obtained with the ALU 247 primers). The meanQ247/Q115 ratio in supernatant samples collected at day 5 and day 7 was 0.50 and 0.38 respectively, suggesting that the DNA released in the supernatant originates mostly from apoptotic rather than necrotic cells.

**B.** Comparison of supernatant-DNA amount when supernatant was stored at RT for different time delays (24h, 48h, 72h and 96h) before extraction. DNA concentrations (pg/ul) were determined using the ALU115 and ALU247 primers.DNA concentration was not affected after supernatant sampling at RT.

**C.** Comparison of supernatant-DNA amount according to the number of freeze–thaw cycles tested on supernatant before extraction. DNA concentration slightly decreased after freeze–thaw cycles applied to supernatantbefore extraction and the quality was not affected.

### Supplemental tables

**Table S1: Articles analyzed to identify genetic abnormalities in hPSC lines.**

