## Supplementary figures and images for "Recurrent genetic abnormalities in human pluripotent stem cells: definition and routine detection in culture supernatant by targeted droplet digital PCR"

### Bubble plot showing recurrent DNA abnormalities

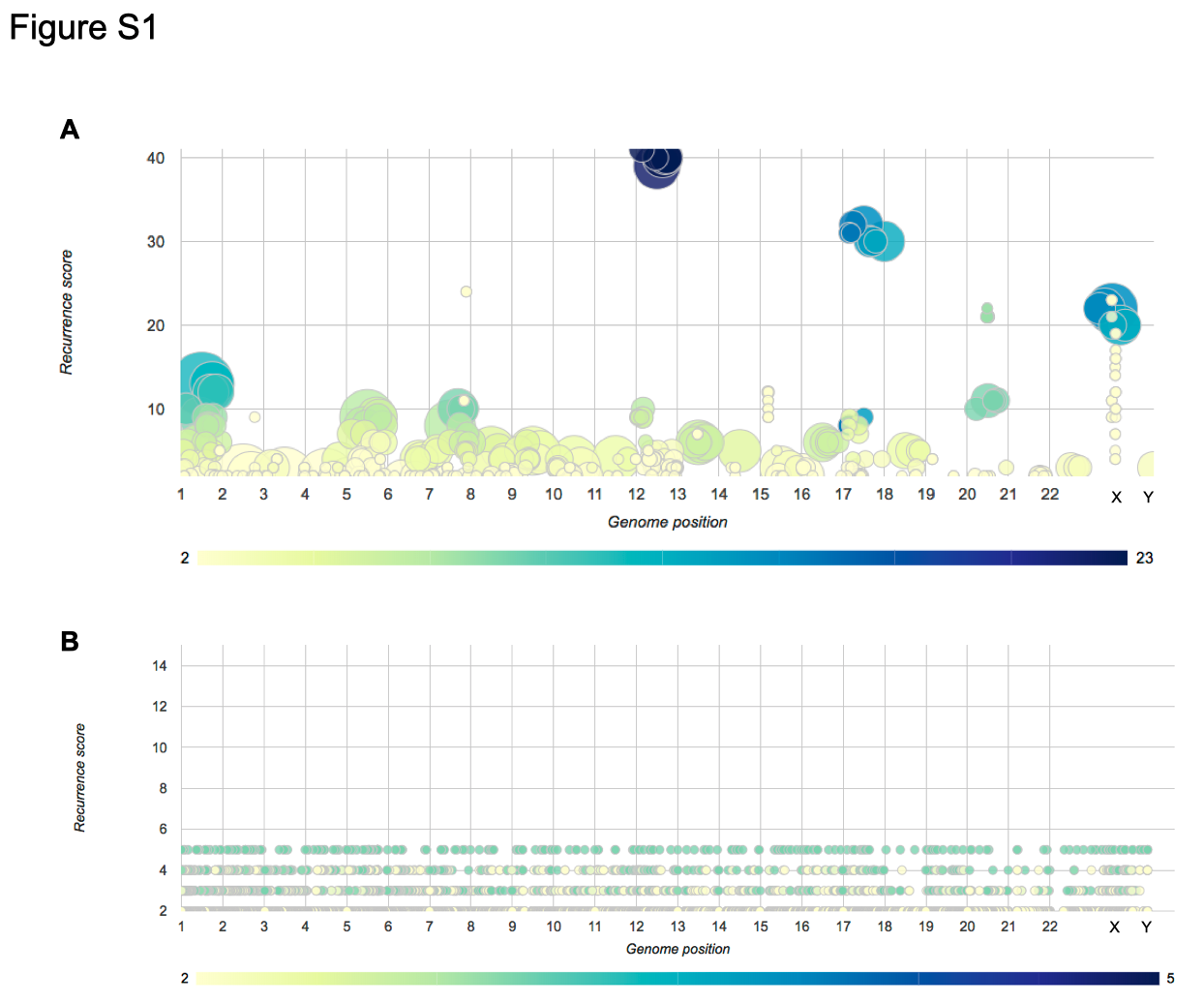

### Effects of various preanalytical conditions on supernatant DNA

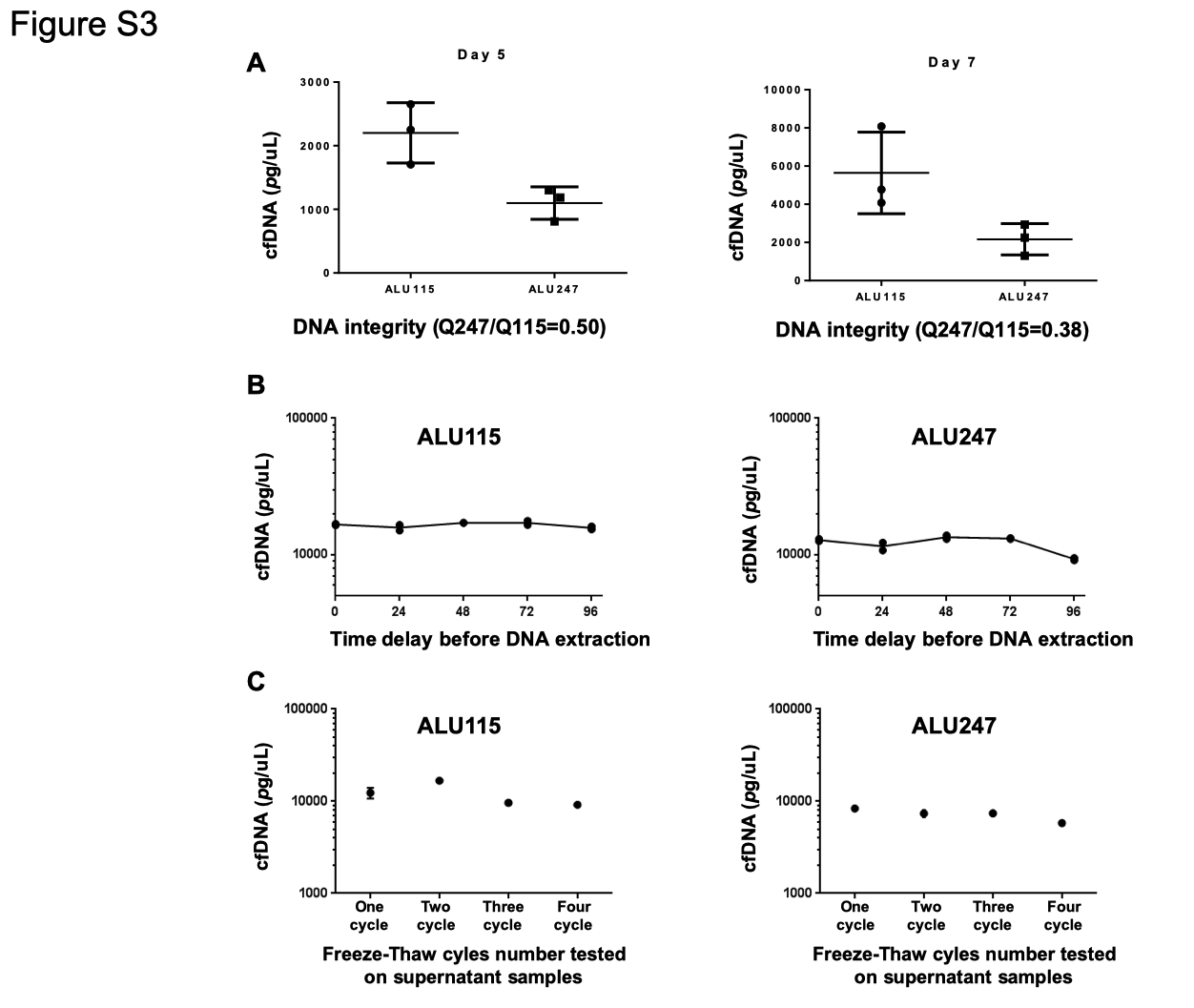

### FACS-like plot illustrating ddPCR results

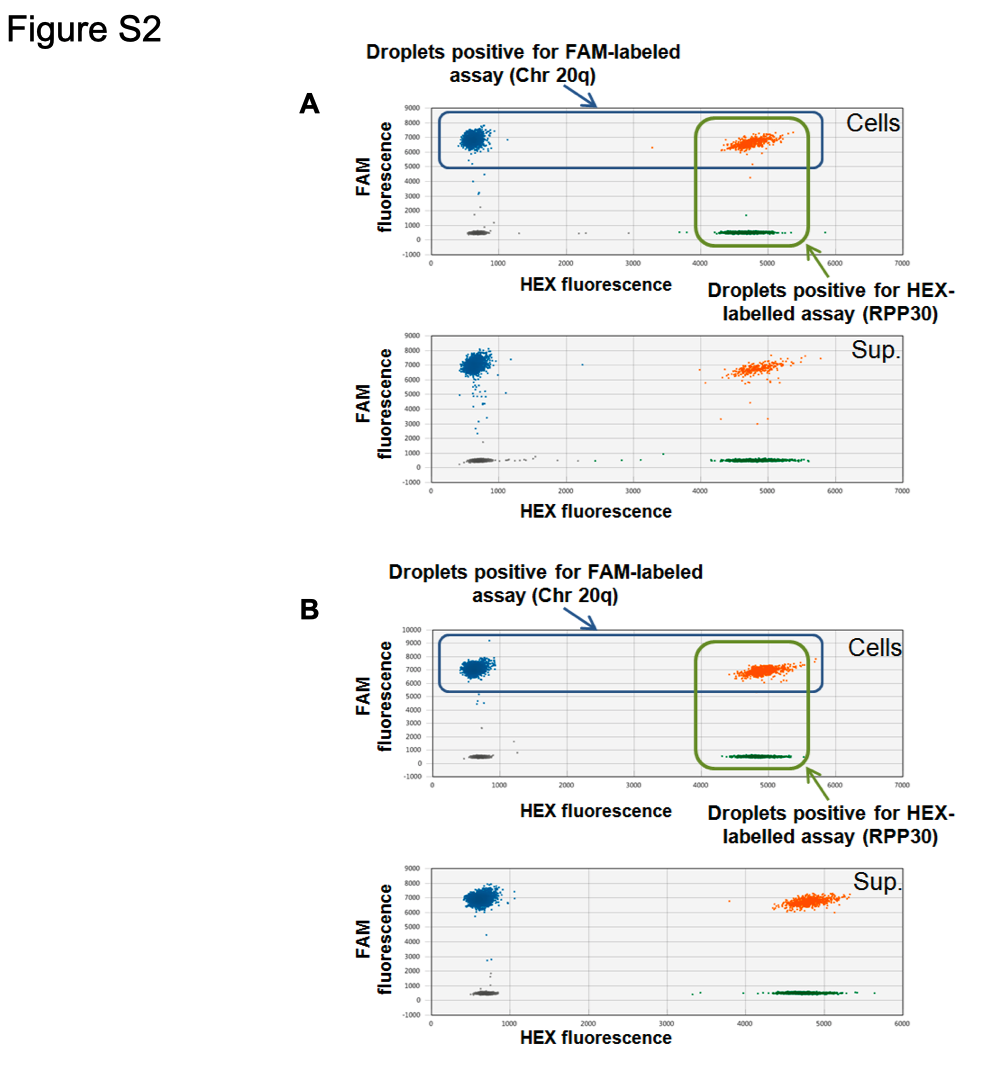
